# Developing Commotio Cordis Injury Metrics by Correlating Chest Force and Rib Deformation to Left Ventricle Strain and Pressure

**DOI:** 10.1101/2020.10.28.359737

**Authors:** Grant J. Dickey, Kewei Bian, Habib R. Khan, Haojie Mao

## Abstract

Commotio cordis is a sudden death mechanism that occurs when the heart is impacted during the repolarization phase of the cardiac cycle. This study aimed to investigate commotio cordis injury metrics by correlating chest force and rib deformation to left ventricle (LV) strain and pressure. We simulated 128 chest impacts using a simulation matrix which included two initial velocities, 16 impact locations spread across the transverse and sagittal plane, and four baseball stiffness levels. Results showed that an initial velocity of 17.88 m/s and an impact location over the LV was the most damaging setting across all possible settings, causing the most considerable LV strain and pressure increases. The impact force metric did not correlate with LV strain and pressure, while rib deformations located over the LV were strongly correlated to LV strain. This leads us to the recommendation of exploring new injury metrics such as the rib deformations we have highlighted for future commotio cordis safety regulations.

## I. INTRODUCTION

Commotio cordis (CC) refers to sudden death from low-energy non-penetrating chest impacts over the cardiac silhouette in the absence of structural heart disease. Defined as a cardiac concussion, commotio cordis shows no signs of structural damage to the heart post-impact (Maron, Poliac, Kaplan, & Mueller, 1995; Pearce, 2005). According to the US Commotio Cordis Registry (USCCR) in Minneapolis, there are currently over 200 confirmed cases worldwide (Link, 2012; Maron & Estes III, 2010). Although the occurrence rate is low, commotio cordis can happen in a wide variety of circumstances; ranging from casual play in a backyard or playground, to competitive hockey, lacrosse, or baseball games (Kaplan, Karofsky, & Volturo, 1993; Maron & Estes III, 2010). Meanwhile, the statistics in cases are believed to be strongly influenced by the lack of awareness towards commotio cordis, suggesting that there are many more cases of the sudden death mechanism that have gone unreported (Maron & Estes III, 2010).

Prevention of commotio cordis has been investigated in the literature and focuses on the use of safety chest protectors. Although the use of chest protectors in contact sports are common, they are not designed with the prevention of CC in mind. One recent study found that a combination of high- and low-density foam, flexible elastomer and a polypropylene polymer in a chest protector reduced the incidence of ventricular fibrillation (VF) by 49% in swine models(Kumar et al., 2017). On the other hand, numerous studies in the literature have explained how commercially available chest protectors fail to reduce the incidence of VF in commotio cordis events(Doerer, Haas, Estes III, Link, & Maron, 2007; Link et al., 2008; Viano, Bir, Cheney, & Janda, 2000; Weinstock et al., 2006). Meanwhile, there is a very specific time window in which an impact to the cardiac silhouette must occur to induce commotio cordis in a subject. The limited-time window makes laboratory investigations challenging, with only a handful of swine experiments with impacts being aimed at the LV successfully causing commotio cordis(Dau, Cavanaugh, Bir, & Link, 2011; Link et al., 1998).

Currently, the National Operating Committee on Standards for Athletic Equipment (NOCSAE) has standard test methods for evaluating chest protectors in their ability to prevent commotio cordis. The evaluation measures peak force over the chest cavity of the NOCSAE Thoracic Surrogate (NTS) through two impact velocity tests (30-mph and 50-mph), with an upper and lower load cell, as well as a cardiac load cell used to measure the impact force. Impact force was measured in newtons (N). For baseballs, peak force from impact in the 30-mph case must not exceed 400 N by the cardiac load cell, and 498 N for the upper or lower load cell. In the 50-mph case, the cardiac load cell must not exceed 800 N, while the upper and lower load cell shall not exceed 1001 N (NOCSAE, 2019). Currently, only force over the chest cavity is included in the testing criteria. Moreover, impact-induced cardiac responses, especially mechanical responses of the LV such as strain and pressure that directly affect the heart remain unknown. Therefore, it is necessary to understand the correlation between external parameters such as chest force and rib deformations to internal heart responses.

This study adopted a detailed finite element (FE) model representing a 10-year-old child chest, which was validated under higher-energy blunt impacts on post-mortem human subjects (PMHSs) (Jiang, Mao, Cao, & Yang, 2013) and was exercised under lower-energy cardiopulmonary resuscitation on live subjects (Jiang, Cao, et al., 2014; Jiang, Mao, Cao, & Yang, 2014). We simulated a total of 128 baseball to chest impacts covering a wide range of real world-relevant events, including various impact velocities, impact locations, and different baseballs. The focus of this study was to understand external forces/deformations to LV strain and pressure. Meanwhile, how different impact settings could affect LV responses were also investigated.

## II. METHODS

### Finite Element Simulation of Baseball to Chest Impact and Post Processing

Impact responses were analyzed using the chest model of the CHARM-10 developed at Wayne State University (Shen et al., 2016), which represents an average 10-year-old child. This detailed FE model includes 742,087 elements and 504,775 nodes. The model contains all major anatomical structures based on detailed clinical scans of 10-year-old children (Mao et al., 2014), including, but not limited to the heart, lungs, chest cavity and kidneys. Another advantage of the chest model is that the model has been validated based on both data collected through cardiopulmonary resuscitation (Jiang, Mao, et al., 2014) on live subjects and impact data collected on cadavers (Jiang et al., 2013). Alongside the validated chest model, a baseball model with a radius of 37.5 mm was created with the material property being defined based on the literature (Dau, 2011).

Each simulation had a run time of 20 ms, with an output frequency of 10,000 Hz. For force, 2,000 Hz for strain and pressure, and 1,000 Hz for deformation. Simulations were run on Ls-Dyna (LSTC, Livermore, Ca). After the simulations were completed, LS-PrePost2.4, an advanced pre and post-processor, was used for data collection and analysis for the impact response (Figure 2).

### Design of Experiments

#### Impact Velocity

The baseball had two initial impact velocities of 13.41 m/s and 17.88 m/s, positioned 1.0 m from the chest cavity (Figure 1. A). These initial velocities are consistent with the literature which reports this as the most susceptible velocity range (Link et al., 2003).

**Figure 1.**
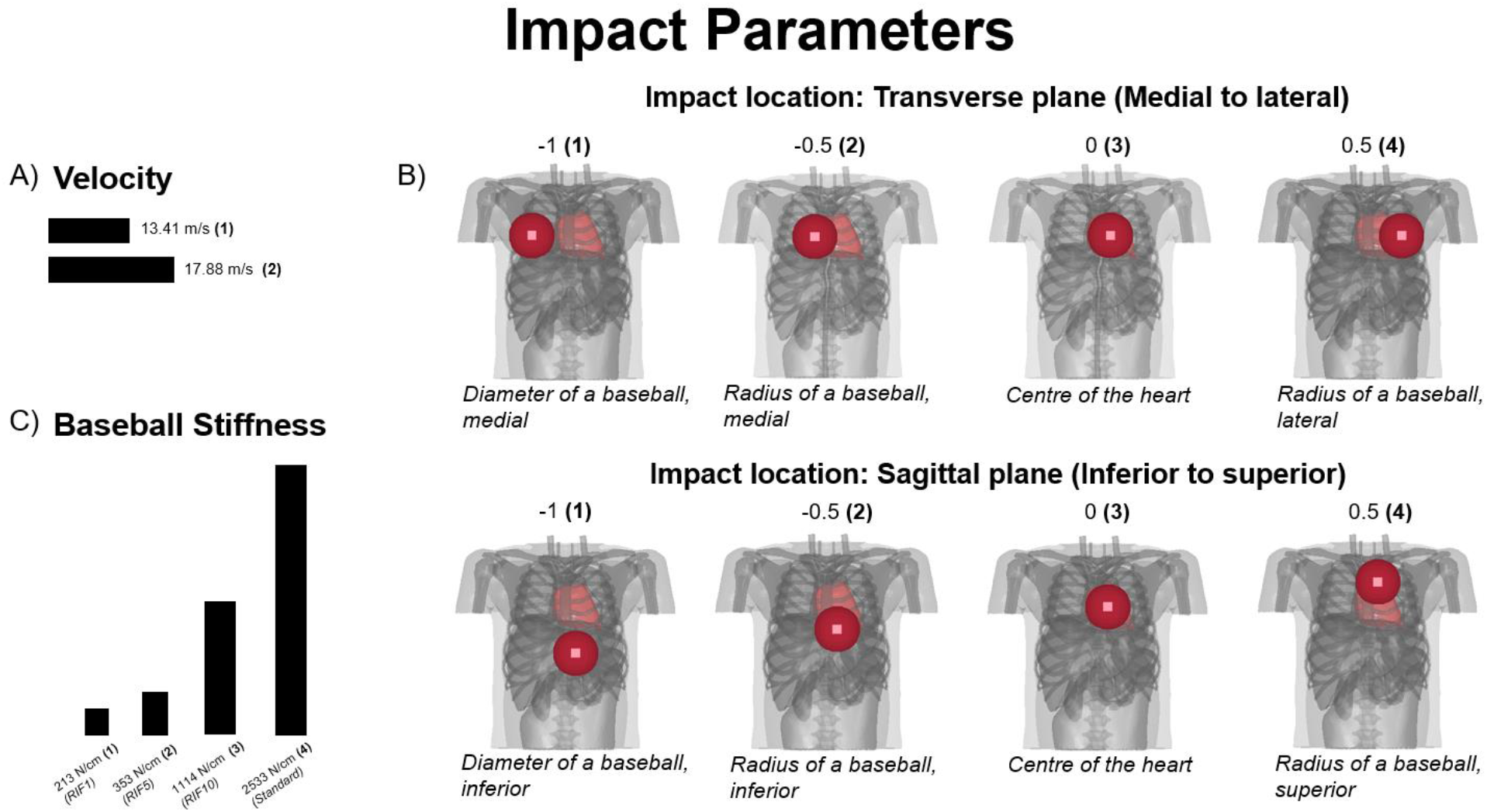
Impact parameters for simulation matrix. A) Impact velocity speeds (m/s). B) Impact locations, transverse and sagittal plane. C) Baseball stiffness levels (N/cm), ranging from soft to a standard baseball.

#### Impact Location

Sixteen impact locations were simulated with the standard direction aiming directly over the heart (Figure 1. B) The baseball moved by the radius of the ball (37.5 mm) medial to lateral (transverse) and/or inferior to superior (sagittal). Together this created four locations in the transverse direction and four in the sagittal direction, for a total of sixteen impact locations.

#### Baseball Stiffness

Four baseball stiffness values were used (Figure 1. C), representing the reduced injury factor (RIF). RIF 1, RIF 5, RIF 10, and standard were simulated. RIF 1 represents a stiffness of 213 N/cm, RIF 5 represents 353 N/cm, RIF 10 represents 1114 N/cm and standard represents a standard baseball stiffness of 2533 N/cm [10].

In total, 128 simulations were conducted, with two impact velocities, sixteen impact locations, and four baseball stiffness values (Figure 1).

#### Impact Responses

Using the CHARM-10 computational model, we analyzed the following impact responses: Force between baseball and chest, max rib deformation, LV strain and pressure, and rib deformation at the LV (Figure 2). Regarding rib deformation at the LV region, we further examined the deformation of rib 3, 4, and 5 as marked in Figure 2.

**Figure 2.**
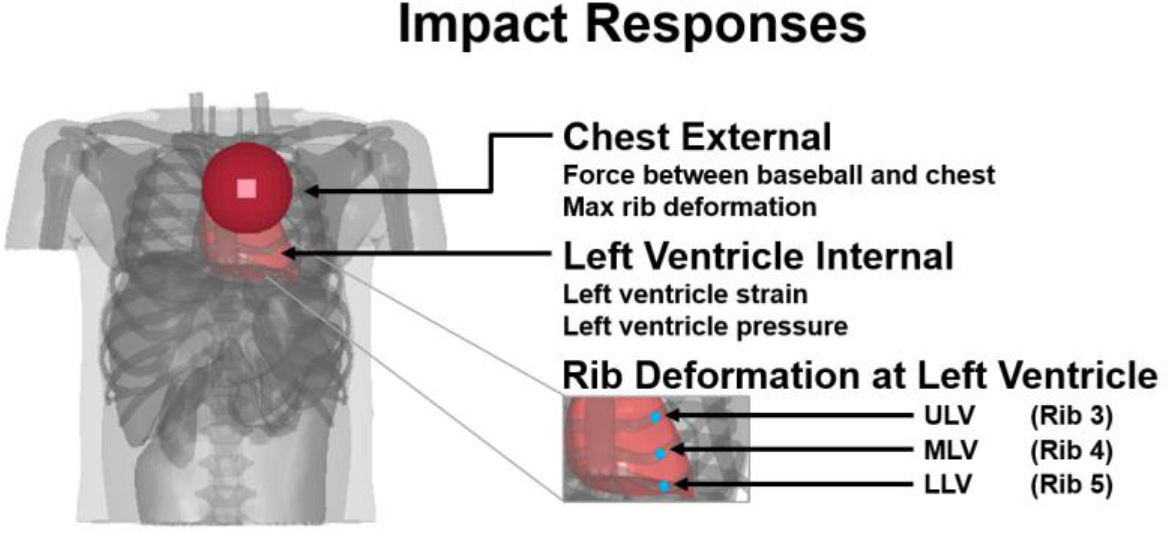
Impact responses analyzed from the computational model. Chest internal responses, left ventricle internal response, and detailed chest external response as rib deformations at the left ventricle region. ULV: Upper left ventricle, MLV: Middle left ventricle, and LLV: Lower left ventricle.

#### Impact Parameter Analysis

Impact parameter results were managed through the use spreadsheets and Minitab (Minitab, LLC, State College, Pennsylvania, USA). The correlation between external response and internal response was conducted through spreadsheet with R^2^ values. Minitab was used to analyze the contribution of each impact parameter by creating a Pareto chart. Main effect charts were created to determine how influential each parameter was in affecting LV strain and pressure.

## III. RESULTS

### Impact Responses vs. Strain Correlations

Both maximum rib deformation and reaction force did not correlate with LV strain with R^2^ values less than 0.01 (Figure 3. A&B). Meanwhile, rib deformation near the upper left ventricle (ULV) and middle left ventricle (MLV) regions did have a strong correlation, with R^2^ of 0.77 and R^2^ of 0.75, respectively (Figure 3. C&D). Rib deformation at the lower left ventricle (LLV) region showed a positive correlation with strain as well, but with R^2^ of 0.34 (Figure 3. E). Overall, ULV and MLV rib deformation correlated with LV strain best (Figure 3. F).

**Figure 3.**
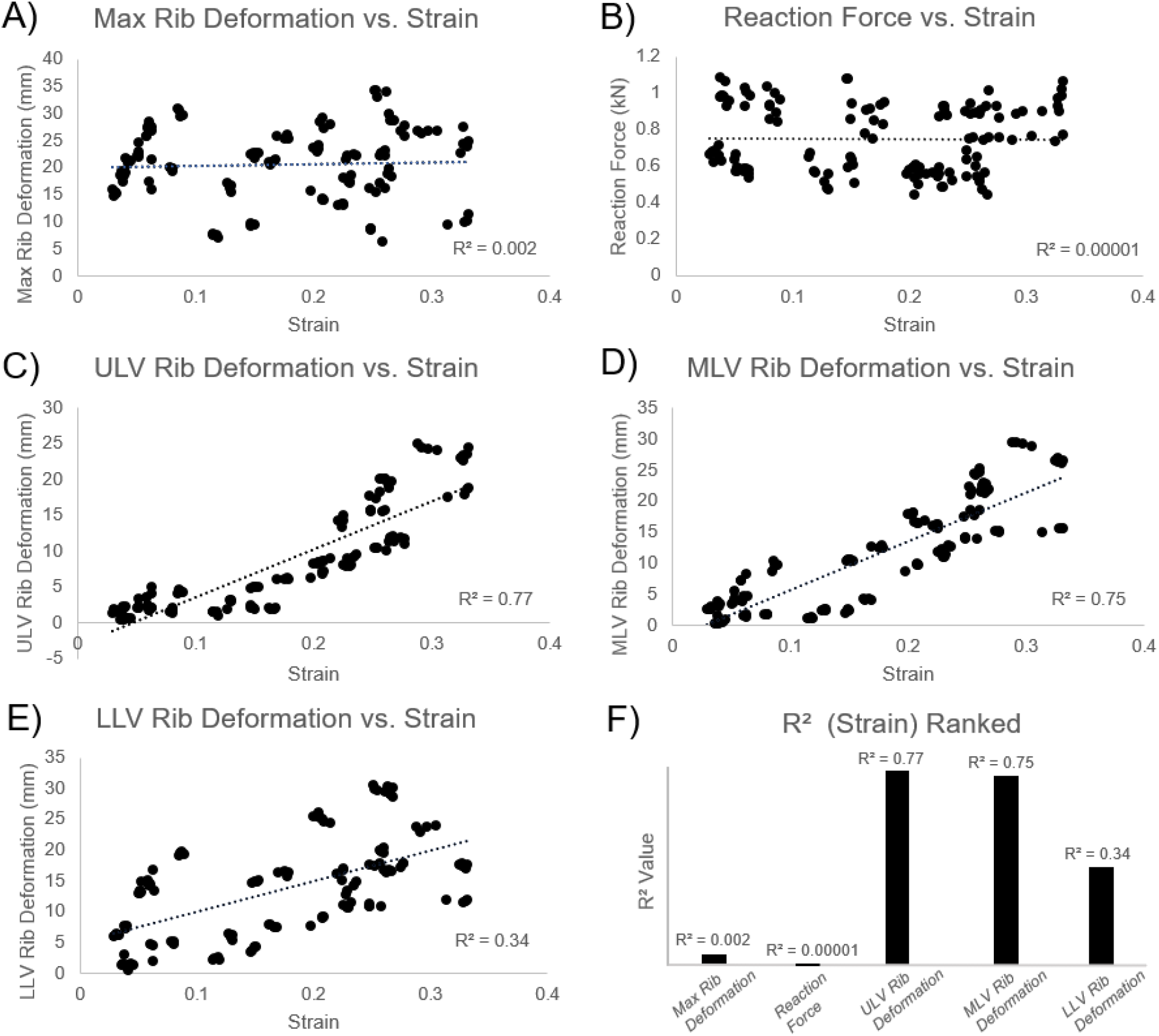
Impact responses vs. strain correlation. ULV and MLV rib deformation have the strongest correlation, with reaction force showing the weakest correlation.

### Impact Responses vs. Pressure Correlations

Similar to LV strain, pressure had a very weak correlation to max rib deformation, and more notably, reaction force (Figure 4. A&B). Reaction force had a very low correlation, with R^2^ values less than 0.1. MLV rib deformation stood out with the strongest correlation and an R^2^ of 0.83 (Figure 4. B), while ULV rib deformation had a strong correlation as well, with an R^2^ value of 0.71 (Figure 4. C). LLV had a moderate correlation with an R^2^ value of 0.52 (Figure 4. E). Overall, MLV and ULV rib deformation correlated with pressure best.

**Figure 4.**
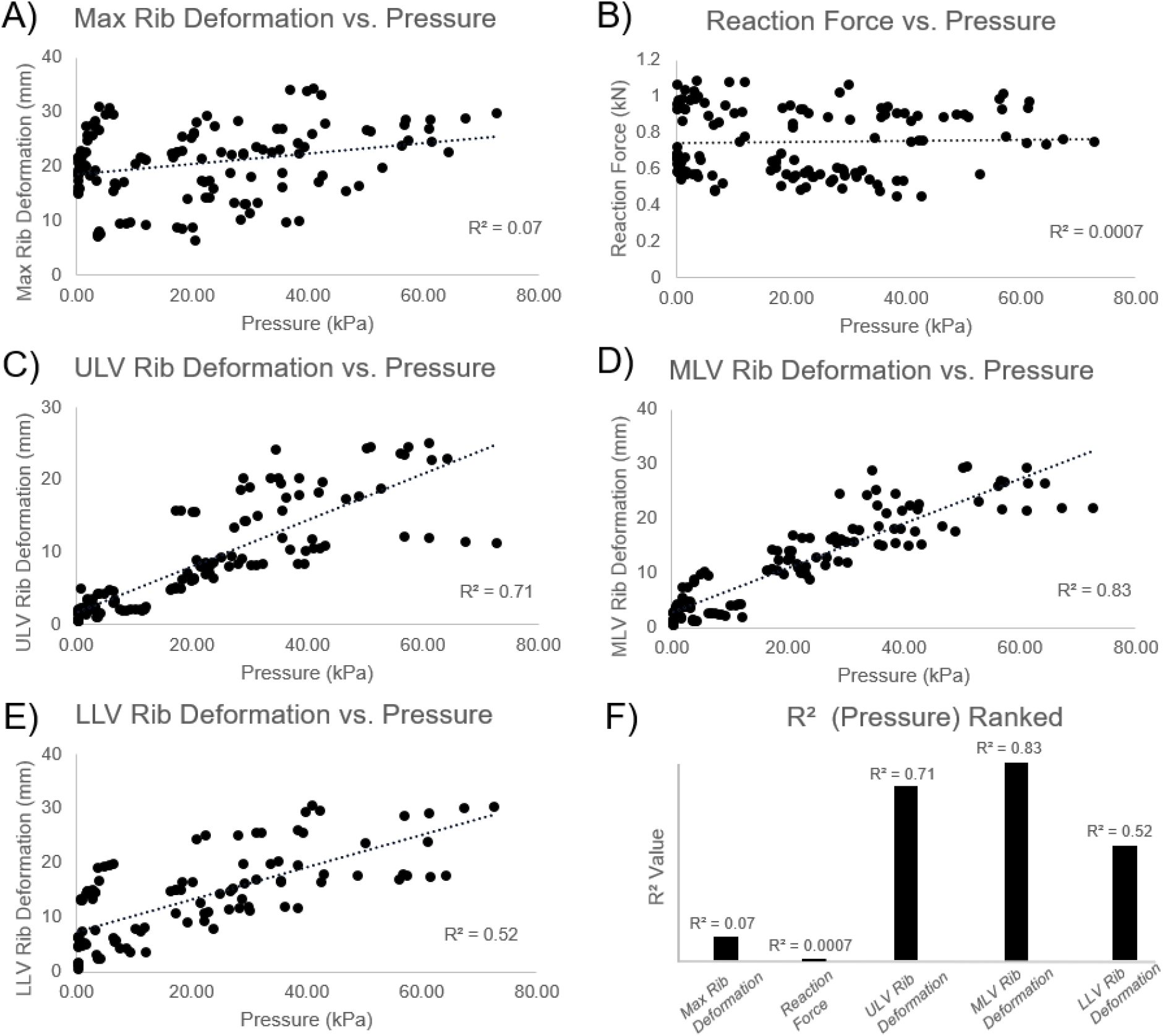
Impact responses vs. pressure correlation. ULVandMLVrib deformation have the strongest correlation, with reaction force showing the weakest correlation.

### Parameters Affecting Left Ventricle Strain and Pressure

Regarding LV strain, the Pareto chart highlighted velocity as the most influential factor (Figure 5. A). Regarding LV pressure, velocity and impact position (transverse and sagittal) are the most important factors (Figure 5. C). Baseball stiffness was found to be an insignificant factor, while changing the baseball from soft to hard stiffness levels did not affect LV strain or pressure (Figure 5. B&D).

**Figure 5.**
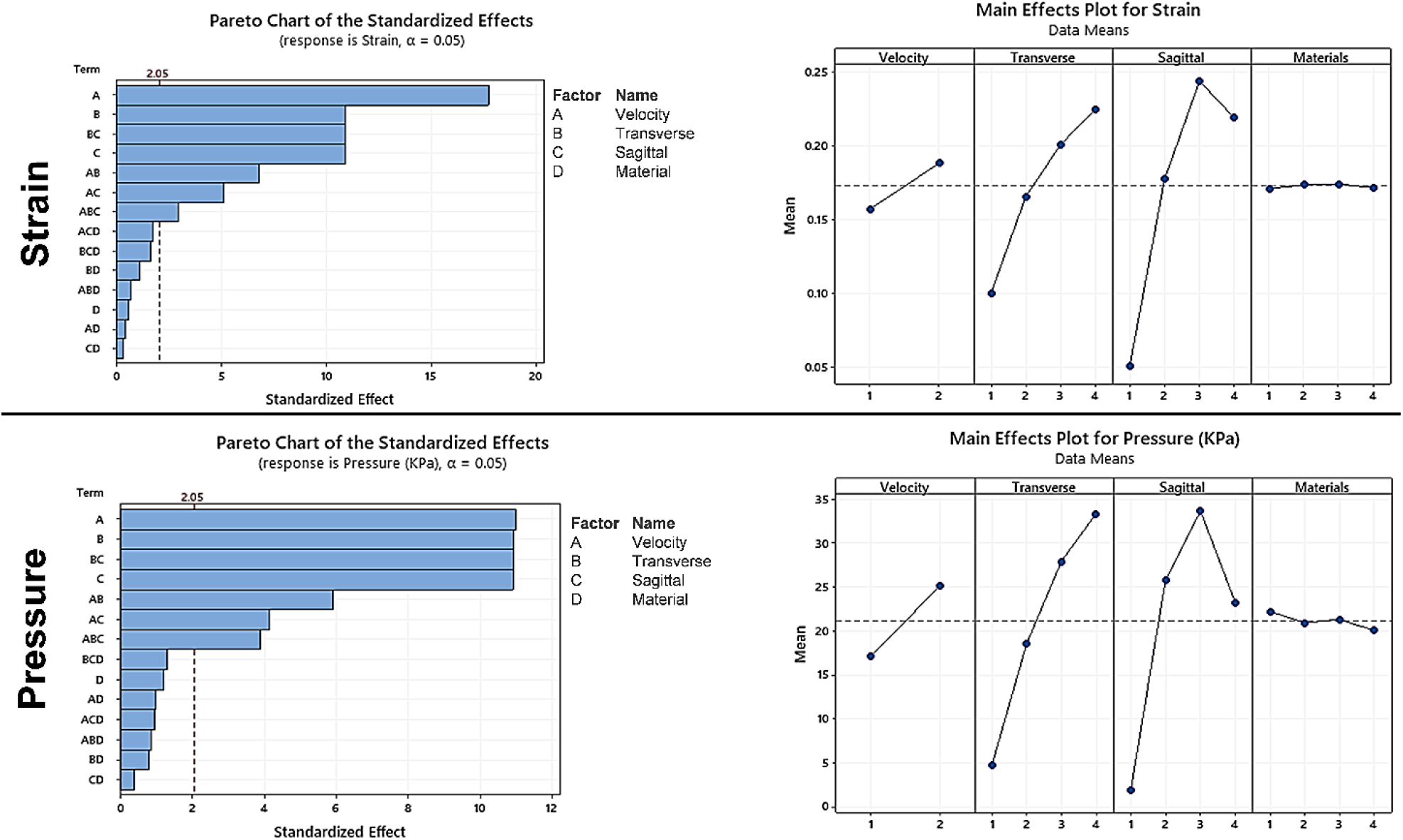
Pareto and main effect charts for strain and pressure of the left ventricle.

### Most Damaging Setting

After analyzing the pareto and main effect charts (Figure 5. B&D), it was concluded that the most damaging setting in all 128 simulations was the combination of an initial velocity of 17.88 m/s and a location in the transverse (4) and sagittal (3) direction (Figure 6). The most damaging setting produced the highest strain and pressure in the LV.

**Figure 6.**
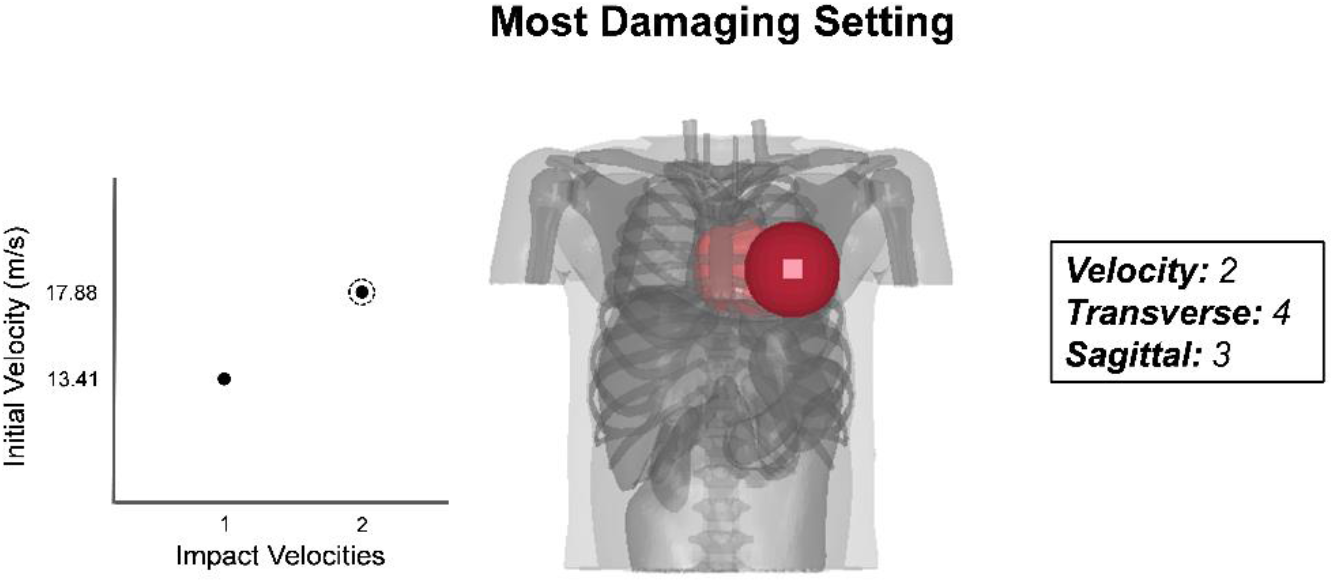
Most damaging setting according to main effect charts. Velocity (2), Transverse (4), Sagittal (3).

### Left Ventricle Strain/Pressure and MLV Rib Deformation Time History

Comparisons were made between two representative high and low velocity cases for LV strain, pressure and MLV rib deformation (Figure 7). Peak strain occurred approximately 5 ms after initial impact, while peak pressure occurred right at the moment of impact. MLV rib deformation peaked around 5 to 10 ms after initial contact, consistent with strain development.

**Figure 7.**
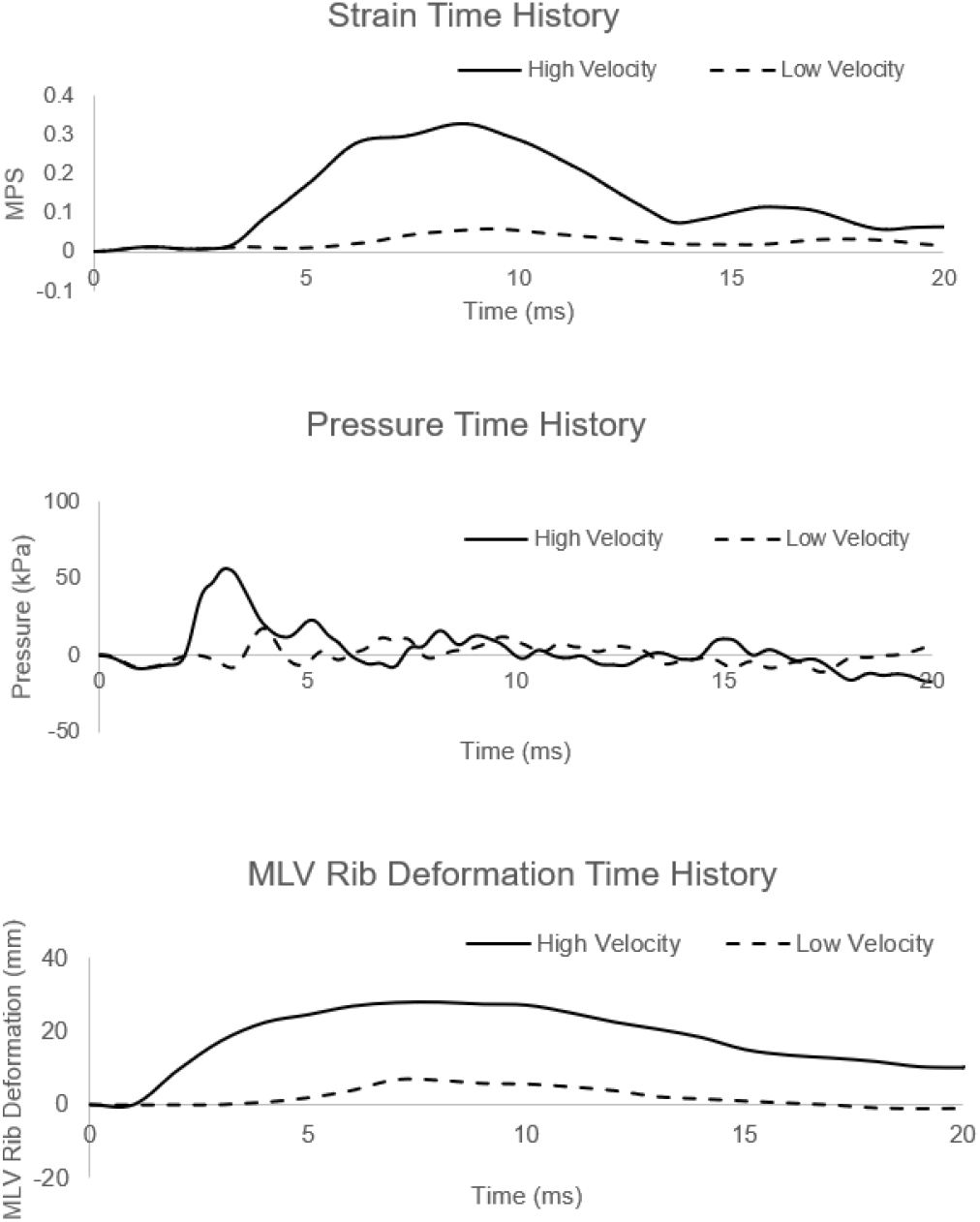
Strain, pressure andMLVrib deformation time history graphs comparing high velocity and low velocity cases. (MPS = Maximum Principal Strain).

### Reaction Force Time History with Filter Comparison

A reaction force time history graph shows the effects of different force filters (Figure 8). The different filter options were selected to further investigate the ability for the NOCSAE accepted low-pass channel frequency class (CFC) 120 filter to collect peak values when looking at reaction force from impacts. We included a no-filter option, as well as a high-pass filter of CFC 1000 for comparisons. Based on the comparison, the filter CFC 1000 was deemed as acceptable and used in this study. Other predictions including strain, pressure, and deformations did not show noise, therefore no filter was applied.

**Figure 8.**
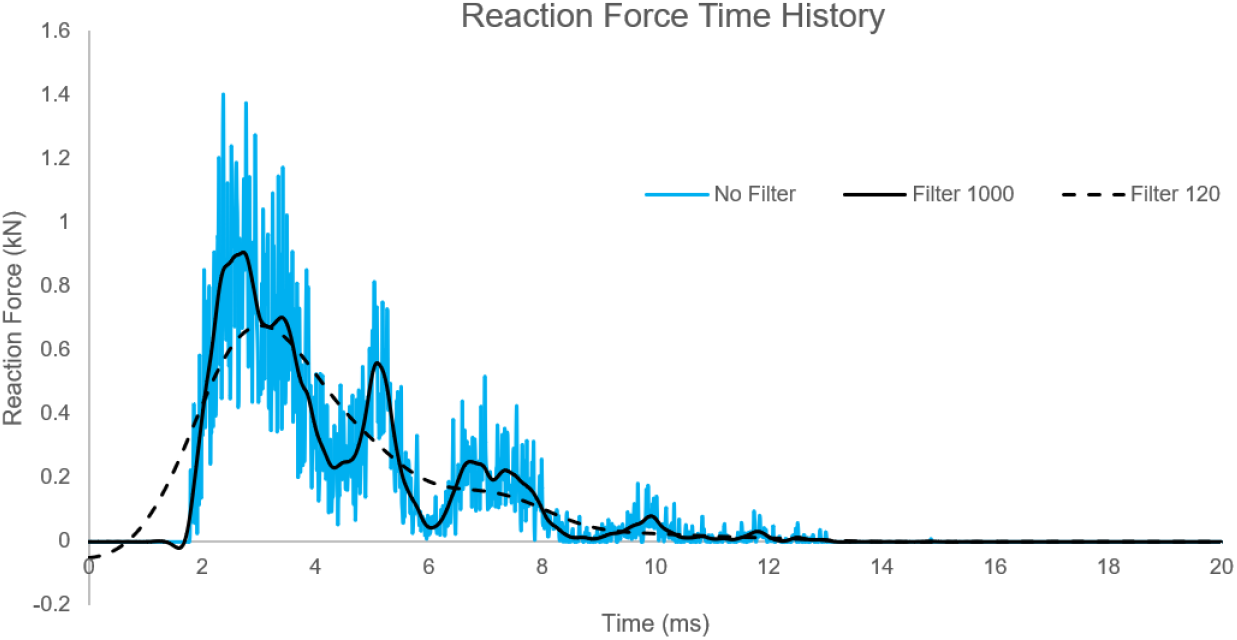
Reaction force time history graph comparing different force filters for one simulation case.

## IV. DISCUSSION

We used a validated child chest model to systematically understand how various chest impacts affected the heart, especially the LV that has been the target in studying commotio cordis. Our data suggested that rib deformation at the upper and middle portion of the LV correlated strongest to LV strain and pressure acutely developed during impacts. However, the impact force or the maximum rib deformation did not correlate with LV strain or pressure, mostly due to varying impact locations. Pareto chart analysis further demonstrated how both impact velocity and impact locations could affect LV strain and pressure. Interestingly, the use of softer baseballs did not reduce LV strain and only slightly reduced LV pressure. To the best of our knowledge, our detailed computational study is the first of its kind to provide data correlating impact parameters and external chest responses to the biomechanical responses of LV strain and pressure using a validated child chest model.

The injury criteria suggested by NOCSAE, as well as current literature, use reaction force to test impact safety (Dau, 2011; NOCSAE, 2019). However, our data shows that there lacks a correlation between impact force and LV strain and pressure (Figure 3, 4). It should be noted that the force sensor specified by NOCSAE was at fixed locations such as the heart position which allows the measurement of force directly applied to the heart region, while the maximum force was measured at the location of impact which could be off from the heart in this study. Considering impacts delivered at various locations, we recommend using ULV and MLV rib deformation as a means of injury metrics for commotio cordis safety, as they both have strong correlations with strain and pressure of the LV.

Initial velocity and impact location are heavily favoured effects in the pareto and main effect charts. Specifically, transverse (4), sagittal (3) and velocity (2) were the highest for LV strain and pressure. Our results from initial velocity match the current literature, a study from 2007 analyzed the effects from different impact velocities on the incidence of VF, finding that a velocity of 40 mph (17.88 m/s) induced VF at a rate of approximately 70%, the highest rate between all impact velocities tested (Madias, Maron, Weinstock, ESTES III, & Link, 2007).

With reference to impact location, our most damaging case depicts an impact over the cardiac silhouette, specifically over the center and base of the LV (Figure 6). Previous studies have identified in swine models that baseball impacts over the LV induce VF during the repolarization phase of the cardiac cycle (Garan, Maron, Wang, ESTES III, & Link, 2005; Link, 2012; Link et al., 2001). Most interestingly, the same study that identified which velocities were most likely to induce VF, reported that it occurred mostly with blows directly over the center of the cardiac silhouette (30% of impacts) compare to those over the LV base (13%) or apex (4%) from a swine model (Madias et al., 2007). It should also be acknowledged that the baseball was moved at 37.5 mm of increments during simulations.

Literature suggests that softer baseballs mitigate commotio cordis events in swine studies (Link et al., 2002). However, our results found that the effect of baseball stiffness levels on LV strain and pressure was not significant, especially when comparing to impact velocities and impact locations. When identifying the effects of varying stiffness levels, we found that for LV strain there was almost no change, whereas for LV pressure there was a mean difference of approximately 1-2 kPa across the 4 stiffness levels. With respect to the Pareto charts, baseball stiffness levels were the clear cut least important factor, showing 95% less significance than initial velocity, and 91% less significant than impact location for left ventricle strain. When considering LV pressure, baseball stiffness was found to be 91% less significant of a factor than initial velocity and impact location. Softer baseballs did slightly reduce LV pressure, which aligns with the recommendation to mitigate commotio cordis events (Classie, Distel, & Borchers, 2010). However, it should be emphasized that chest protection is greatly needed even with using softer baseballs.

We used the 1000 Hz filter rather than the 120 Hz filter specified in the NOCSAE standard test methods for commotio cordis protectors (NOCSAE, 2019). The reason we chose this filter was because the low-pass filter 120 may have been too strong for our data, therefore missing higher peak values of data and ultimately missing high reaction force values. The low-pass 120 filter had an 18% decrease in its’ ability to measure peak force values when compared to a low-pass 1000filter. The 120 filter measured a peak of 0.65 kN, while the 1000 filter measured 0.90 kN. Meanwhile, it should be acknowledged that the filter 1000 was deemed as acceptable for out computational prediction which is different from the experimental force sensors.

Although our model is exceedingly detailed, one of our limitations is that the model does not include fluid structures inside the heart. Further studies could attempt to incorporate blood flow through the heart to simulate a more realistic simulation with more computational power being available to solve the fluid-structure interaction with deformable boundaries. Nevertheless, we have applied an internal pressure of 9.3 kPa to the heart wall mimicking blood pressure and focused on collecting acute strain and pressure raised in milliseconds during the impact, which was expected to not be affected by the flow change in this short period of duration.

## V. CONCLUSION

This detailed computational study helps to address the tissue-level biomechanical mechanisms of commotio cordis. Initial velocity and impact location in the transverse and sagittal plane over the chest cavity played important roles in LV strain and pressure, whereas baseball stiffness made almost no difference in this regard. Left ventricle strain and pressure correlated strongly with ULV and MLV rib deformation, while they did not correlate with maximum reaction force from the baseball to chest. From our understanding of the current literature and through this study, we should place a more considerable emphasis on the heart’s response during and after initial impact and develop injury metrics based on this knowledge. Impact responses such as strain and pressure of the LV and deformation of the ribs located at ULV and MLV could prove to be appropriate measurements in the evaluation of chest protectors through commotio cordis testing methods.

## VI. ACKNOWLEDGEMENTS

We acknowledge the support of NSERC and the Canada Research Chairs program.

